# Endoplasmic Reticulum morphological regulation by RTN4/NOGO modulates neuronal regeneration by curbing luminal transport

**DOI:** 10.1101/2021.05.10.441946

**Authors:** Tasuku Konno, Pierre Parutto, David M. D. Bailey, Valentina Davì, Cécile Crapart, Mosab Ali Awadelkareem, Colin Hockings, Aidan Brown, Katherine M. Xiang, Anamika Agrawal, Joseph E. Chambers, Molly Vander Werp, Katherine Koning, Emmanouil Metzakopian, Laura Westrate, Elena Koslover, Edward Avezov

## Abstract

Cell and tissue functions rely on an elaborate intracellular transport system responsible for distributing bioactive molecules with high spatiotemporal accuracy. The tubular network of the Endoplasmic Reticulum (ER) constitutes a system for the delivery of luminal solutes it stores, including Ca^2+^, across the cell periphery. The physical nature and factors underlying the ER’s functioning as a fluidics system are unclear. Using an improved ER transport visualisation methodology combined with optogenetic Ca^2+^ dynamics imaging, we observed that ER luminal transport is modulated by natural ER tubule narrowing and dilation, directly proportional to the amount of an ER membrane morphogen, Reticulon 4 (RTN4). Consequently, the ER morphoregulatory effect of RTN4 defines ER’s capacity for peripheral Ca^2+^ delivery and thus controls axonogenesis. Excess RTN4 limited ER luminal transport, Ca^2+^ release and iPSC-derived cortical neurons’ axonal extension, while RTN4 elimination reversed the effects.

**Summary:** Intracellular transport through the lumen of the ER network is modulated through narrowing/dilation of ER tubules by a membrane morphogen – RTN4, a process controlling axonogenesis by limiting the delivery of ER-stored Ca^2+^.

## Results and discussion

The Endoplasmic Reticulum (ER) is the single largest organelle, forming stacks of membranous sheets contiguously interconnected with a network of tubules that extends throughout the cell periphery. This architecture enables the ER to deliver its unique contents (e.g. Ca^2+^) throughout the cell and to supply numerous contact points with other organelles (e.g. mitochondria, endo/lysosomes). While macro-fluidic distribution systems on the organism level are well understood, the physical nature and structure-function relationship of the ER nano-fluidics network remains obscure. In terms of ER morphogenesis, considerable advances have helped elucidate how membranes can be curved to attain the tubular ER structure. In particular, a family of ER shaping proteins has been identified and substantially characterised^1^. Independently, a neuron-specific repertoire of these ER morphogens has emerged, and an increasing number of members are associated with neurodegenerative diseases both biochemically^2–5^ and as hereditary causes^6–9^. Remarkably, two ER shaping protein-variants (of approximately two dozen known examples) were sufficient to reconstitute the ER structure *in vitro*^10^. This suggests a potential regulatory function, other than simple morphogenesis, for at least some of the remaining non-essential morphogens such as RTN4. This gene is a relatively late evolutionary acquisition (appearing with the advent of amphibians), dispensable for cell survival and morphogenesis of ER tubules^11–13^. Earlier studies designated RTN4 as Nogo reflecting its axonal plasticity-restricting properties^14^. An excess of Nogo/RTN4 splice variant-a limits axonal regeneration^15^, while its knockout improves the process in animal models^11,12^. Thus, we sought keys to unravel how modulating ER structure may regulate the organelle and the neuronal cell function, by delineating the effects of RTN4 on ER structure-function.

To assess the effect of RTN4 on tubular ER ultra-structure, we visualised the endogenous protein in sub-diffraction resolution along with an ER luminal marker (Fig. 1A). In agreement with previous studies, the images showed a heterogeneous distribution of the protein across the network^16^ (Fig. 1A, Video 1). Notably, the RTN4-enriched tubular areas showed a profound reduction in the luminal marker’s signal (Fig. 1A, B & Video 1). The observed luminal content exclusion by RTN4 suggested that the morphogen may constrict ER tubules (Fig. 1A and B). This notion is supported by the fact that RTN4’s Reticulon Homology Domain (RHD) domain, responsible for its ER membrane localisation, assumes a wedge-like configuration that can impose curvature on the tubule membrane^1,17^. Consistently, another reticulon showed a similar effect when overexpressed^18^. To test whether and to what extent RTN4 can modulate ER tubular diameter, we measured its changes in response to RTN4 overexpression and knockout. The scale of ER tubule cross-section (<100 nm^19^) is below the resolution limit of light microscopy. To surmount this limitation, we extract the tubular diameter by analysing fluorescent intensities of an ER membrane marker, a technique compatible with live-cell imaging. It draws on the principle that the membrane marker signal intensity on a cylindrical structure (ie: ER tubule) is proportional to its radius, while the intensity on a flat structure (ER sheet) depends only on its area. Thus, tube radius R was approximated by equation (1):

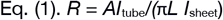

where *I*_sheet_ is signal intensity on a peripheral sheet of area A, and *I*_*tube*_ the intensity along proximal tubules of length *L*. These measurements yielded values consistent with previous estimates^19,20^ Fig. 1C). In relative terms, these measurements confirmed a dose-dependent narrowing of the ER tubules by exogenous RTN4 (Fig. 1C). Remarkably, the elimination of RTN4 expression by CRISPR/Cas9 knockout measurably increased the ER diameter (Fig. 1D, Fig. S1A & B), consistent with EM estimates in fixed embryonic fibroblasts of RTN4 KO mouse^21^. Eliminating RTN from human neuroblastoma cell line (SH-SY5Y) led to a more conspicuous increase in tubule width, suggesting that endogenous RTN4 burden in higher in these cells, compared to COS-7 (Fig. 1D).

**Figure 1.**
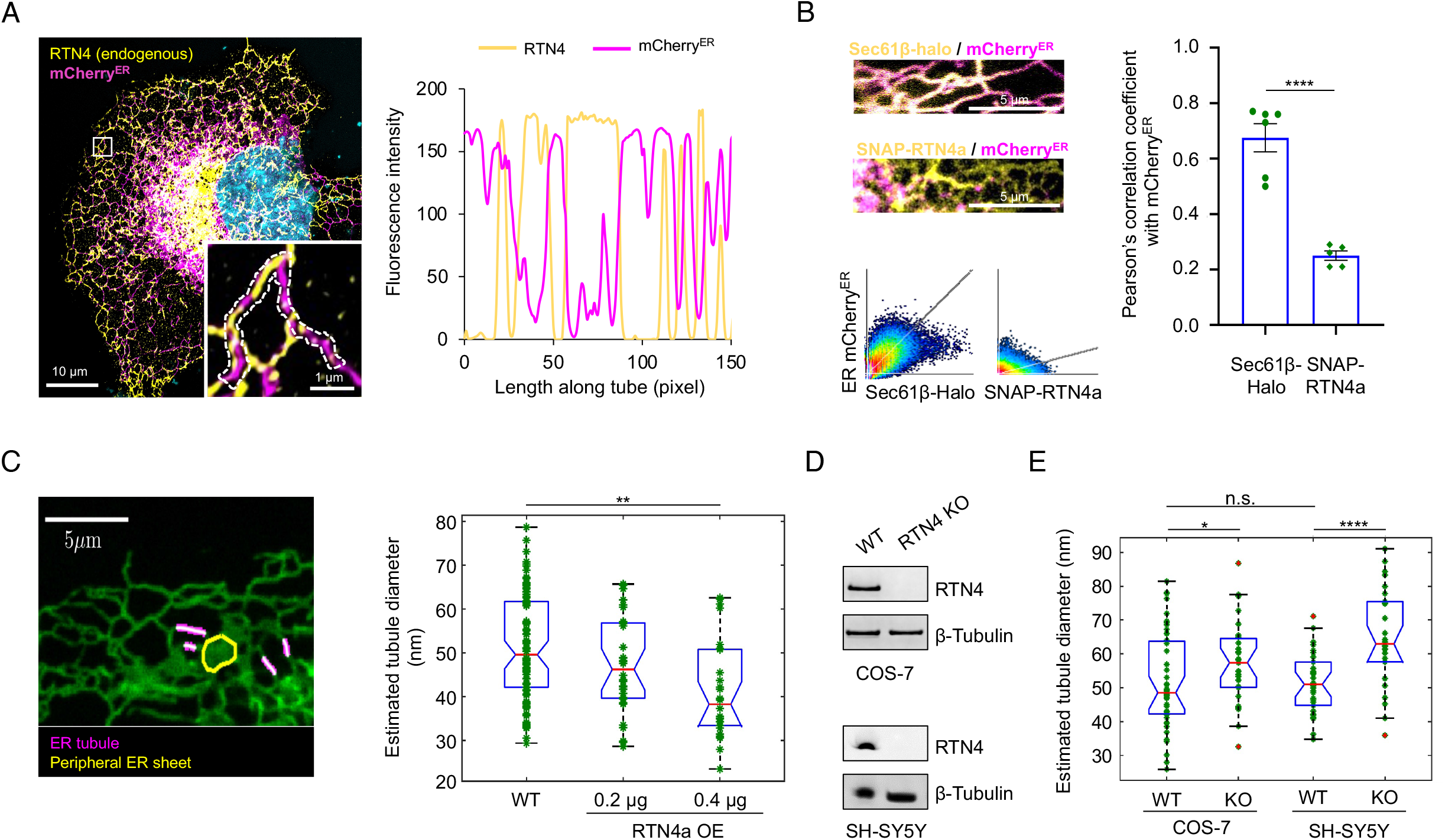
RTN4 excess narrows ER tubules. A. Fluorescence micrographs of endogenous RTN4 (immunolabeled) and an exogenously expressed ER luminal marker (mCherry^ER^) in fixed COS-7 cells, shown along with a line-scan analysis of the fluorescence intensity along peripheral ER tubules (dashed line in inset). B. Colocalisation analyses of the ER luminal (mCherry^ER^) vs overexpressed membrane (Halo-Sec61β or SNAP-RTN4a) markers. Note the high colocalisation of mCherry^ER^ with Sec61β but not with RTN4a. C. Estimated ER tubule diameter in COS-7 cells, transfected with increasing amounts of RTN4a, co-expressed with ER membrane markers (seem methods for details). D. Immunoblot of RTN4 in normal and CRISPR knockout (RTN4 KO) COS-7 and SH-SY5Y cells. E. Measurements as in C of RTN4 KO cell lines. (COS-7 WT; n=47, COS-7 RTN4 KO; n=26, SHSY5Y WT; n=30, SHSY5Y RTN4 KO; n=26). \**P* < 0.05, \*\**P* < 0.01, \*\*\**P* < 0.005, \*\*\*\**P* < 0.001, n.s., not significant).

We hypothesised that this observed alteration of ER tubular diameter by RTN4 may modulate luminal content transport through the ER network and, consequently, control the rate of material exchange across long cellular distances. To test this hypothesis, we measure intra-organelle transport by tracing the spatiotemporal distribution of a locally photo-injected membrane or luminal photoactivatable fluorescent protein^22^ (Fig. 2A-D & S2A, Intensity PhotoActivation Chase, iPAC). Namely, we assessed the timescale of photoactivated signal arrival at increasing distances from its origin – the cell area continuously exposed to UV laser illumination, photo-activating the tracer-proteins (Fig. 2A & Video 2). The arrival kinetics are reflected in the effective half-time for signal rise at each local region of interest (intensity time traces in Fig. 2A & B). As visualised by a physical simulation (Fig. 2C & Video 3), the dependence of the signal rise time on distance from the photo-injection origin provides a measure of transport dynamics and elucidates the mode of motion. Namely, diffusive transport corresponds to a quadratic scaling of arrival time with distance from the origin (Fig. 2B & D red curves), as would be expected for a system with diffusive particles spreading from a region of fixed concentration^23^. By contrast, transport on an active network with locally processive motion yields signal rise times that scale less than quadratically with distance (Fig. 2B & D blue curves). In agreement with this calculation, measurement of the signal spread kinetics for an ER membrane and luminal markers showed a diffusive-like behaviour for the former and evidence of faster super-diffusive transport for the latter (Fig. 2B). This observation mirrors the active flow model of ER luminal transport^24^. Notably, paGFP^ER^ also showed a pulsatile pattern of photoactivated signal spread across the network (Video S2). Although the mechanism of flow generation in the ER remains to be established, this observation supports a pumping-like effect of contraction-relaxation of ER structure elements (tubules or junctions^24^. Thus, the iPAC methodology provides a sensitive means to quantify transport behaviour across a range of cellular distances.

**Fig 2.**
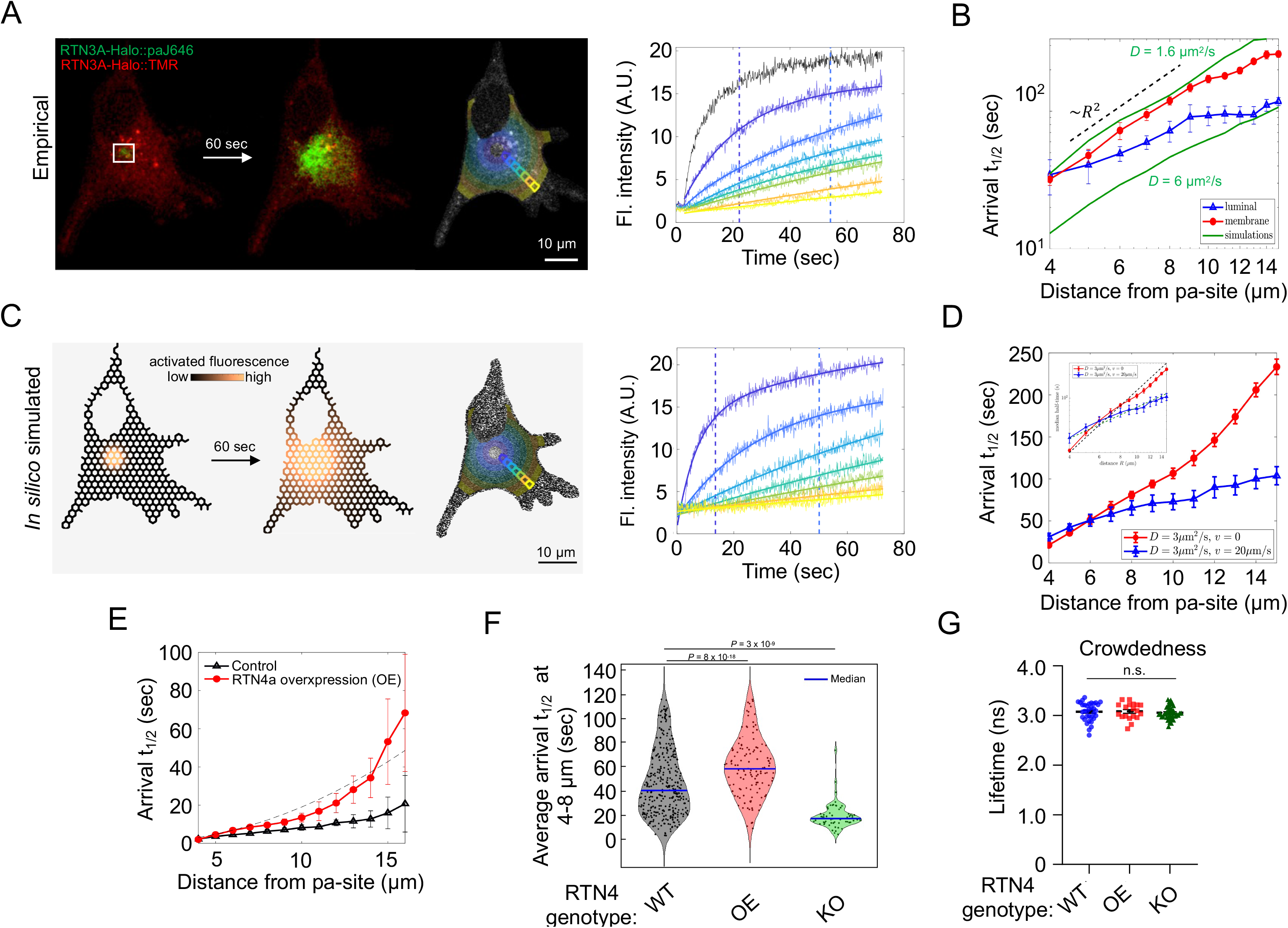
ER morphoregulation by RTN4 modulates ER luminal transport rates. A. Representative images of intensity photo-activation chase (iPAC) in COS-7 cells transiently expressing RTN3A-Halotag, labelled by tetramethyl rhodamine (TMR) and photoactivatable-Janelia Fluor 646 (paJ646). White box denotes the area of paJ646 photoactivation by laser illumination 405 nm. Traces of photoactivated signal intensity are coloured according to the distance from the photoactivation spot. B. Median halftime of photoactivated signal-rise (arrival t_1/2_, exemplified by vertical dashed lines in A) for ER-targeted photoactivatable GFP (paGFP^ER^) or an ER membrane marker (Sec61β-Halo::paJ646), at various distances from the activation spot. Note, measurements for membrane protein mirror simulations of diffusive spread at D = 1.8 μm^2^/s. Luminal protein measurements show transport with super-diffusional scaling and cannot be fit by higher diffusion coefficients (compared to simulations up to D = 6 μm^2^/s, green lines). C. *In-silico* molecular motion model, simulating diffusive transport of proteins from photoactivation region in the ER tubular network (for a diffusion coefficient, *D* = 3 μm^2^/s). Simulation results processed identically to experimental data (in A and B), generating curves of local concentration over time, are shown on the right. D. The simulated median arrival t_1/2_ plots at various distances from the origin as in B, comparing spread by exclusively by diffusion (red) and active network (blue), which includes randomly oriented flows of velocity (*v* = 20 μm/s) in each edge, persisting over τ = 0.1 sec. Note, active flows result in a superdiffusive (sub-quadratic) scaling of signal arrival times versus distance (inset). E. paGFP^ER^ arrival time measurements as in B in mCherry^ER^ (Control) or mCherry-RTN4a overexpressing (RTN4 OE) COS-7 cells (n = 15 each) and F. in RTN4 OE and KO SH-SY5Y cells (*P* derived from Kolmogorov-Smirnov test). G. Fluorescence lifetime (FLT) measurements of an ER-localised molecular crowding probe in cells as in F (n = 34, 19, 42, respectively, n.s., not significant).

We applied iPAC to characterise how RTN4-mediated ER tubule narrowing affects transport rates throughout the ER. Modulation of active luminal flows provides a plausible mechanism for coupling ER tubule diameter with spatial transport rates. The classic Poiseuille flow model for pressure-driven motion of a viscous fluid through a narrow tube predicts that the flow speed *v* scales with radius *R* according to (*v* ∼ *R*^2^)^25^. Furthermore, a lower radius could reduce the persistence time of flows that result from tubule contraction. Accordingly, iPAC revealed that overexpression of RTN4 conspicuously slowed the arrival of locally photoactivated paGFP^ER^ at increasing distances (Fig. 2E & Fig. S2B automated and manual analyses, accordingly. In contrast, RTN4 knockout cells showed a detectable increase in protein mobility through the ER network, as reflected in faster arrival of photoactivated signal at distant regions (Fig 2F & S2C). Thus, the changes in protein mobility, observed upon manipulations of RTN4 expression (Fig. 2E & F) mirror their tubule width-modulating effect: RTN4-imposed narrowing of ER tubules correlates with slower luminal material transport, whilst tubule widening in RTN4 knockout appears to improve material exchange across the ER network (without a detectable RTN4 effect on luminal crowdedness, Fig. 2G & S3). It should be noted that a simple decrease in tubule radius, without modulating transport velocities, would merely rescale the fluorescence intensity measurements throughout the network and would not be expected to alter photoactivated signal rise times.

Next, we set out to examine the consequences of transport speed modulation on ER function. We postulated that ER morphological alteration affects its performance as a Ca^2+^ supply system. Namely, RTN4 overexpression is expected to reduce Ca^2+^ delivery within the ER through two complementary effects – the reduced flux arising directly from a lower cross-sectional area, and the reduced long-distance transport speeds as observed in iPAC measurements. This is supported by a physical simulation of Ca^2+^ transport across the ER network and its ability to supply local releases (Fig. 3A & Video 4). The simulation incorporated diffusive and active transport of both free Ca^2+^ and its binding buffer proteins on a realistic ER network structure, focusing on the total amount of the ion delivered to a release region. It predicts that both ER tubule narrowing and attenuation of processive transport will diminish Ca^2+^ release rates, with the combined effect of a substantial decrease in Ca^2+^ release capacity (Fig 3A).

**Fig 3.**
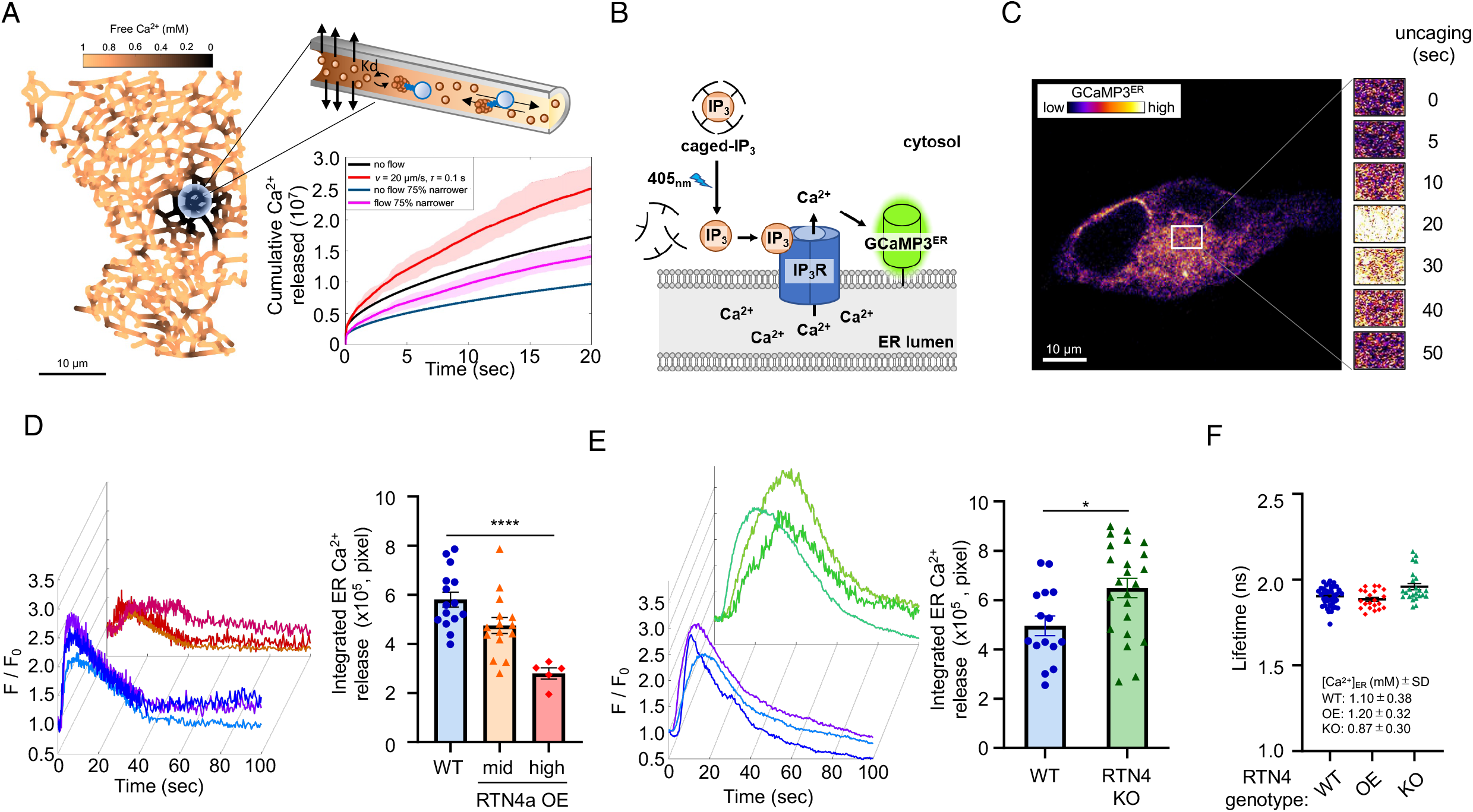
RTN4 ER morphoregulatory/transport effect curbs the ER Ca^2+^ release capacity. A. Physical simulation of the dependence between ER Ca^2+^ release capacity and luminal transport, incorporating equilibrated binding to Ca^2+^ buffer proteins, local ER release, diffusive luminal transport of free/buffered Ca^2+^ and luminal flow, with plots of total Ca^2+^ released over time. Note, release capacity decreases both as a result of halting active flows and from the direct decrease in flow rate due to tubule narrowing. B. Schema of light-induced ER Ca^2+^ release and monitoring assay. C. Representative fluorescence intensity image series of GCaMP3^ER^ photo-uncaging regions of COS-7 cell preloaded with caged-IP_3_ (3 μM, 3 hours, uncaging by continuous 405 nm laser illumination). D. Traces of GCaMP3^ER^ signal as in C in WT or RTN4a overexpressing (RTN4a OE) SH-SY5Y cells, shown along with the integrated ER Ca^2+^ released during uncaging period (the area under the curve, WT; n = 17, RTN4a OE; n = 20). Show are means ± SEM from samples in three independent experiments. \*\*\*\**P* < 0.001 (one-way ANOVA). E. ER Ca^2+^ release measurements as in D. for WT and RTN4 KO SH-SY5Y cells (WT; n = 15, RTN4 KO; n = 23). \**P* < 0.05 (student’s *t* test). F. Fluorescence lifetime imaging microscopy (FLIM) measurements of ER Ca^2+^ using the D4ER probe in cells as in E & D. Note Lifetime is inversely proportional to [Ca^2+^].

To test this prediction in a regime that mimics physiological Ca^2+^ release, we used an ER membrane-tethered sensor (GCaMP3^ER^) to track changes in cytoplasmic Ca^2+^ in response to a local photo-lysis-release of inositol-trisphosphate (IP_3_, Fig. 3B & C). IP_3_ is generated in natural signalling as a transient second messenger that triggers ER Ca^2+^ release by binding the ER-membrane Ca^2+^ channel - IP3-receptor^26,27^. In this modality, Ca^2+^ release was dampened by RTN4a overexpression, compared to normal cells (Fig. 3D, amount of Ca^2+^ release reflected in the area under the curve). In contrast, RTN4 knockout cells showed a more robust release capacity (Fig. 3E, no significant decrease/increase in ER Ca^2+^ content was measurable in RNT4 OE/KO to explain the above effect Fig. 3F, S4). The RTN4-dependent changes in the ER’s ability to release Ca^2+^ mirror the effect of this morphogen on tubule diameter and transport (Fig. 1 & 2). Thus, in agreement with the model (Fig. 3A), RTN4-mediated decrease of ER luminal mobility led to weakened Ca^2+^ release.

Given the crucial role of ER Ca^2+^ release at the leading edge of outgrowing/ regenerating cellular projections, including neurites^28^, the effect of RTN4 on this process may explain its plasticity-restricting (Nogo) effect. Accordingly, RTN4-mediated attenuation of the ER’s capacity to fuel Ca^2+^ releases may limit neurite outgrowth (axonal plasticity/regeneration). To test this hypothesis, we assessed the effect of RTN4 on neurite outgrowth. To exclude the possibility of an inhibitory effect from RTN4 on the surface of oligodendrocytes (a possibility previously suggested based on known regeneration pathways^29^, we experimented in oligodendrocyte-free cortical neuronal monocultures, which we derived from human induced Pluripotent Stem Cells (iPSC^30^, Fig. 4A). In these neuro-physiologically active cells (Video 5 & Fig. S5), both endogenous and overexpressed RTN4a localised exclusively to the ER (Fig. 4B & C). Conspicuously, exogenous RTN4 inhibits neurite outgrowth as the majority of RTN4 overexpressing cells failed to generate neurites (Fig. 4D & E). Though viable, RTN4a-overexpressing cells did not show neurite outgrowth during differentiation (Fig. 4E). Further, neurons not undergoing plasticity events tolerated RTN4 in their neurites’ ER when overexpression was imposed post-differentiation (Fig. 4F). Conversely, RTN4 KO improved neuronal outgrowth during iPSC differentiation (Fig. 4D, with no effect on total cell mass, S1C, 6, & Video 6). In agreement with animal model studies^11^, axonal regeneration was also improved by RTN4 knockout, but in this case, the effect occurred in an oligodendrocyte-free neuronal monoculture with exclusively ER-localised RTN4 (Fig. 4G, H & Video 7).

**Fig 4.**
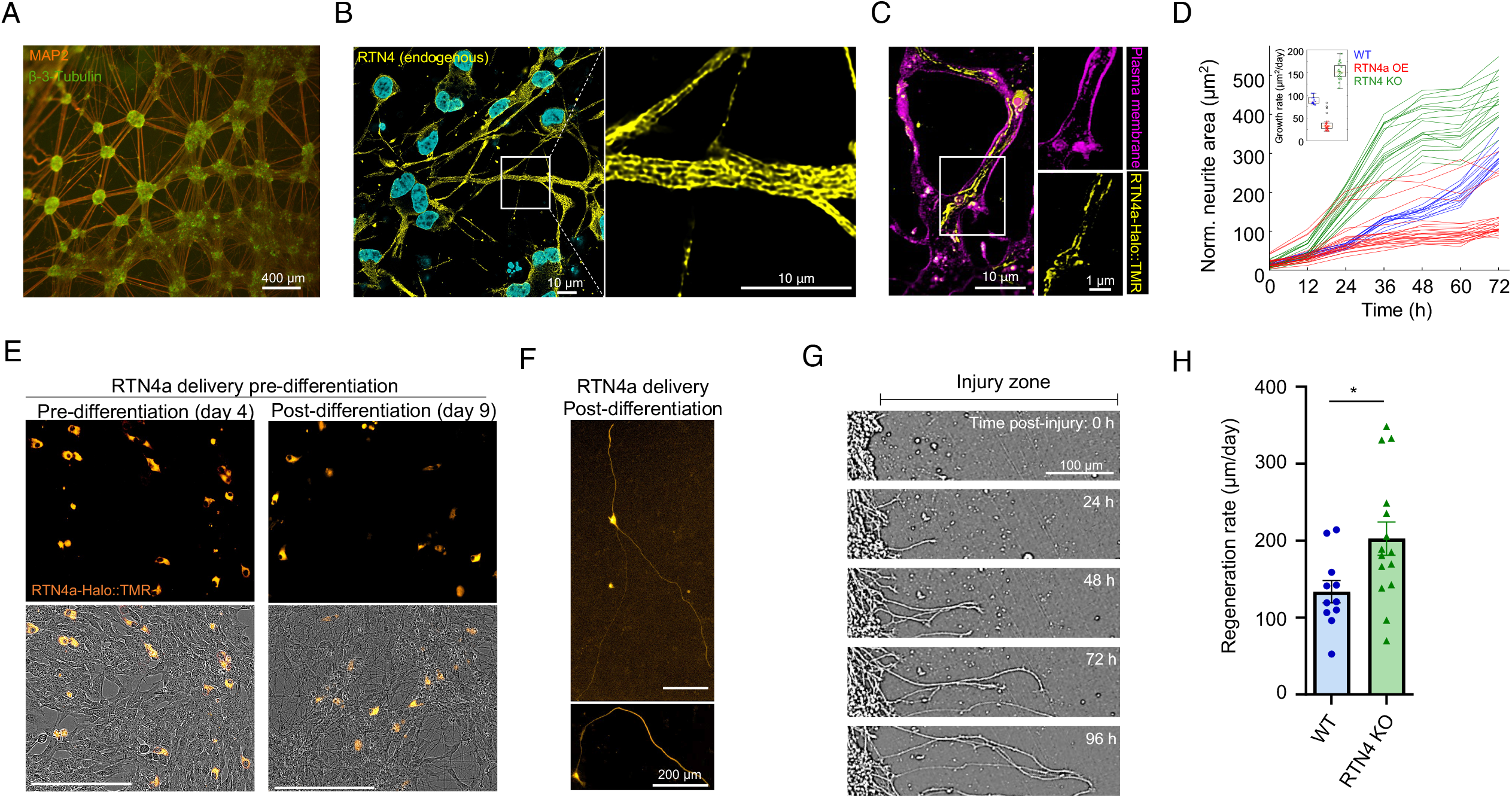
RTN4 hinders neuronal outgrowth/regeneration in human iPSC-derived cortical neurons. A. Representative image of differentiated human iPSC-derived cortical neurons (iNeurons) stained by neuron-specific marker MAP2 and β-3-Tubulin. B. Micrographs of immunolabelled endogenous RTN4 in fixed iNeurons with sub-diffraction-limit resolution (nuclei label - Hoechst 33258), and C. of exogenous RTN4a (RTN4a-Halo::TMR) co-stained with a plasma membrane marker (Cellbright). D. Normalised area covered by neurites during outgrowth of WT, RTN4a overexpressing (OE) or knockout (KO) iNeurons, inset: corresponding growth rate extracted from a linear fit. E. Micrographs of iNeurons with TMR-labelled RTN4a-Halo (orange) at the indicated cortical neuron differentiation stage. F. As in E, but in this case RTN4a-Halo was introduced post-differentiation (Day 14). G. Representative time-lapse images of neurite regeneration following a mechanical injury. H. Neurite regeneration rate in WT and RTN4 KO iNeurons. Shown are means ± SEM from three independent experiments.

Taken together, these findings indicate that ER morphoregulation can modulate material distribution throughout the cell periphery, tuning Ca^2+^ release profiles. The endogenous level of RTN4 expression is sufficient to substantially constrain ER transport, as RTN4 knockout leads to improved transport kinetics (Fig. 2) and stronger Ca^2+^ release (Fig. 3). A key functional consequence of ER morphoregulation by RTN4 is the modulation of axonal plasticity (Fig 5). This result highlights a hitherto unknown coupling between ER structure and its function as an intracellular delivery system, specifically in neurons. This mechanism may exemplify a physiological regulatory principle operating on the nanoscale in analogy to the vasoconstriction/vasodilation of body’s circulation system.

**Fig 5.**
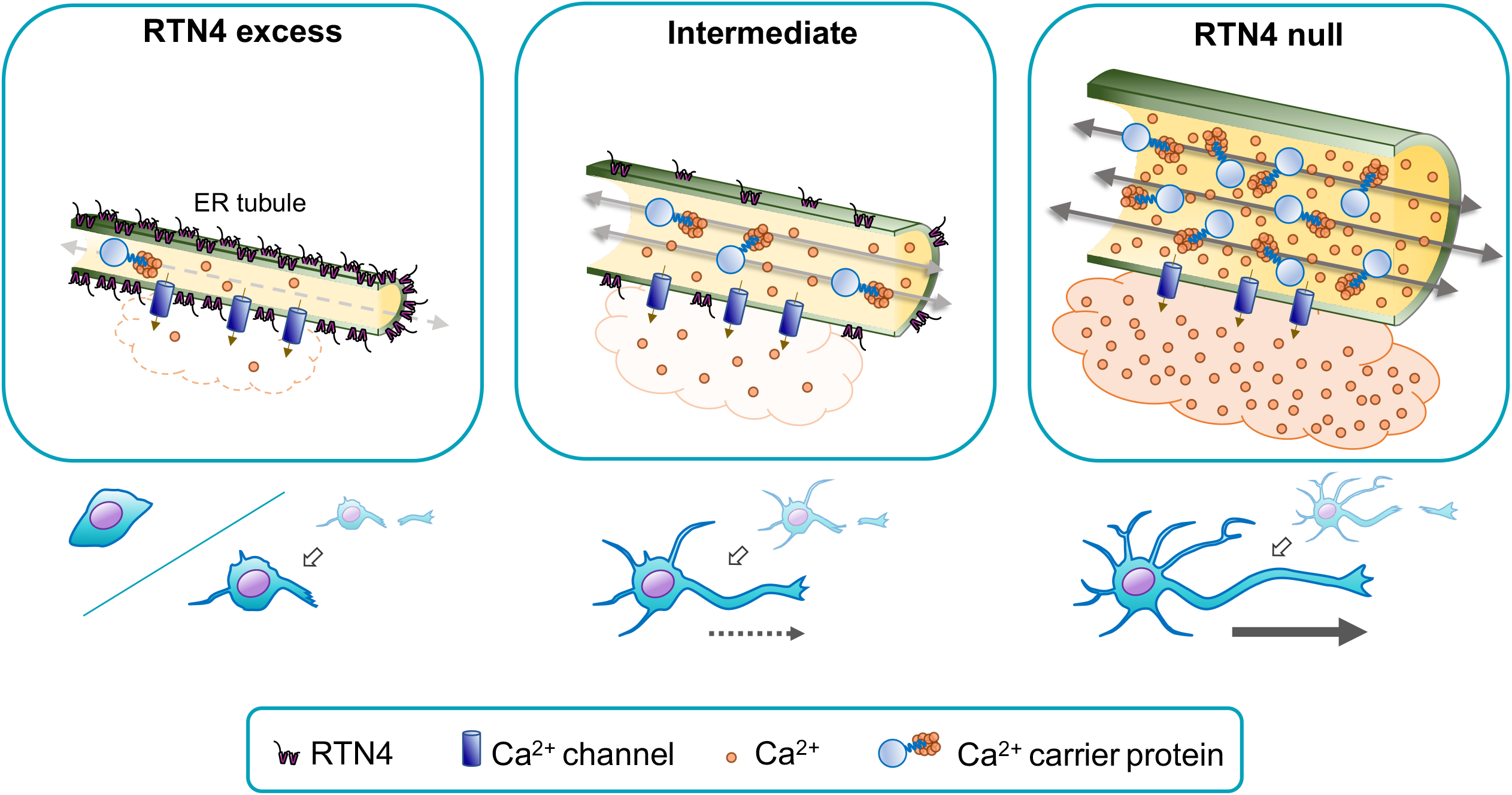
Schematic of RTN4-mediated modulation of ER luminal transport with an effect on Ca^2+^ distribution and consequently axonal outgrowth/regeneration.

## Acknowledgements

We are extremely grateful to Luke D. Lavis (HHMI Janelia Research Campus, USA) for providing multiple Halo-tag fluorescent ligands including photoactivatable-Janelia Fluor 646 and to Gregory Strachan (Institute of Metabolic Science, University of Cambridge, UK) for invaluable technical supports for setting up photoactivatable pulse-chase assays. This work is supported by grants for the UK Dementia Research Institute, which receives its funding from UK DRI Ltd, funded by the UK Medical Research Council, Alzheimer’s Society and Alzheimer’s Research UK and Alzheimer’s Society grant (AS-595) to EA. EFK and LMW were supported by the US National Science Foundation Grant #2034482. Additional support for EFK was provided by a Cottrell Scholar Award from the Research Corporation for Science Advancement and by the Hellman Fellows Fund.

## Methods

### Cell culture, transfections, and expression constructs

COS-7 (RRID:CVCL_0224) and HEK293T (RRID:CVCL_0063) cells were cultured in Dulbecco’s modified Eagle’s medium (DMEM) supplemented with 10% fetal bovine serum (FBS), 2 mM L-Glutamine and 100 U/mL Penicillin-Streptomycin (P/S). SH-SY5Y (RRID:CVCL_0019) cells were cultured in DMEM/F12 (1:1) medium supplemented with 10% FBS, 2 mM L-Glutamine, 100 U/mL P/S and 1 × non-essential amino acids (NEAA, 11140035, Gibco). Transfections were performed using the Neon Transfection System (Invitrogen). Descriptions of the plasmids used in this study were presented in Supplemental Table 1.

### Lentiviral preparations

Lentiviral particle was produced by transfecting HEK293T cells with the RTN4a-Halo expression vector and three helper plasmids expressing Gag, Pol, Rev, and VSVG. The transfection was carried out using the Polyethylenimine (PEI) with a plasmid ratio of 1:1:1:1. The lentivirus-containing medium was harvested 48 and 72 hours after transfection and pre-cleaned with a 3,500 g centrifugation and a 0.45 μm filtration. The lentivirus-containing medium was subsequently collected by an ultracentrifugation (100,000 g for 2 hours). After a removal of the supernatant, the pellet of lentiviral particle was re-suspended in PBS.

### Human iPS cells culture and differentiation

Human iPSCs with Neurogenin-2 (NGN2) transgene stably integrated into a ‘safe-harbour’ locus under a doxycycline (Dox)-inducible promoter were cultured in TeSR-E8 medium (05990, STEMCELL Technologies). Differentiation into iPSCs-derived cortical neurons (i^3^Neurons) was performed according to previously described protocols with slight modifications ^31^. Briefly, at day 1 and day 2 of differentiation, iPSCs were cultured in DMEM/F12 medium supplemented with 1 x N-2 supplement, 2 mM L-Glutamine, 1 x NEAA, 50 nM 2-Mercaptoethanol (2ME), 100 U/mL P/S, and µg/mL Dox and changed daily. After day 3, culture medium was replaced to Neurobasal medium (Thermo Fisher) supplemented with 1 x B-27 supplement, 2 mM L-Glutamine, 50 nM 2ME, 100 U/mL P/S, and 1 µg/mL Dox, 10 ng/mL NT-3, and 10 ng/mL Brain-derived neurotrophic factor (BDNF). Full-media changes made up until day 6, after which, half-media changes were made every other day.

### Antibodies and reagents

Rabbit polyclonal antibodies to RTN4 (ab47085) and chicken polyclonal antibodies to MAP2 (ab5392) were obtained from Abcam. Mouse monoclonal antibodies to beta-tubulin (MA5-16308), Alexa Fluor 488-conjugated goat anti-rabbit secondary antibodies (A-11034), Alexa Fluor 555-conjugated goat anti-mouse secondary antibodies (A-21137), and Alexa Fluor 568-conjugated goat anti-chicken secondary antibodies (A-11041) were obtained from Invitrogen. Caged-IP_3_ (ci-IP3/PM, #6210) was obtained from Tocris. HaloTag TMR (G8251) and Janelia Fluor 646 HaloTag Ligand (GA1120) were obtained from Promega (Madison, WI, USA). SNAP-Cell 505-Star (S9103S) and SNAP-Cell 647-SiR (NEB S9102S) were obtained from New England BioLabs (NEB, Ipswich, MA, USA). Thapsigargin (10522-1mg-CAY) was obtained from Cambridge Bioscience (Cambridge, UK).

### Immunofluorescence analysis

Prior to immunofluorescence staining, cells were fixed with 2% PFA, 2% glutaraldehyde, 100 mM cacodylate (pH 7.4), and 2 mM CaCl_2_ for 30 min at room temperature (RT), permeabilised with 0.5% Triton X-100/PBS for 30 min at RT, and blocked with 10% goat serum/PBS for 30 min at RT. Primary antibodies were used with 1:1000 dilution for overnight at 4 °C and secondary antibodies were used with 1:1000 dilution for 1 hour at RT. Images were acquired with a confocal microscope (STELLARIS8, Leica, Wetzlar, Germany) with lightning deconvolution.

### Microscopy – imaging and analyses

#### Quantifying tube diameter from relative fluorescence analysis

To calculate the average radius of ER tubules, we leveraged the integrated fluorescence intensities from labeled ER membrane proteins (Calnexin in Fig. 1C and Sec61β in Fig. 1D). Calnexin images were obtained as follows. COS-7 cells (ATCC) were grown at 37°C with 5% CO_2_ in Dulbecco’s modified Eagle’s medium (Gibco) containing 10% fetal bovine serum and 1% penicillin/streptomycin (Invitrogen). Emerald-Calnexin was a gift from Michael Davidson (Addgene plasmid #54021) and mCherry-RTN4a was previously described ^32^. Plasmid transfections were performed as described previously ^33^ and the following standard amounts of DNA were transfected per ml: 0.2 µg Emerald-Calnexin with either 0.2/0.4 µg mcherry_RTN4a or 0.2/0.4 µg mCherry_empty as a control (see figure for respective concentration and transfection setup). Twenty-four hours post transfection, cells were fixed at room temperature with 4% paraformaldehyde plus 0.5% glutaraldehyde in PBS. All images were acquired on an inverted fluorescence microscope (Nikon A1 confocal microscope) equipped with a 100x NA 1.45 oil objective.

We assume that the labeled protein concentrations per membrane surface area are constant throughout a given image, and that the integrated fluorescence intensity in any cellular region is therefore proportional to the total ER membrane surface area in that region. For a region containing a single tubule of radius *R* and length *L*, the total fluorescence intensity is taken as *I*_tube_ = *a(2πRL)*, where *a* is a prefactor incorporating protein density and fluorescent intensity per protein. For a peripheral sheet region, assumed to correspond to a single flat ER structure, surrounded by two membranes of area *A*, the integrated fluorescence intensity is taken as *I*_sheet_ = *a(2A)*. By comparing intensities in a peripheral sheet and nearby tubule regions, the tubule radius can be approximated from the ratio of the two, according to the relationship in Eq. 1

To process each individual image, background intensity is determined by selecting a region outside of the ER network and taking the median intensity in that region. The background signal is subtracted from the image prior to analysis. We then manually identify a peripheral sheet region with well-defined nearby tubules (eg: Fig. 1C). We selected peripheral sheets, rather than those near the nucleus, to avoid challenges associated with stacked sheet structures. To avoid edge effects, the sheet ROI is eroded by a disc structuring element of radius 0.2 µm. Nearby tubule regions are selected such that no tubule intersections or nearby encroaching tube or sheet regions appear alongside each individual tube segment. To incorporate fluorescence from the diffraction-limited area around each tube segment, the backbone of each segment is dilated by 0.2 µm, yielding the pink regions in Fig. 1C. The length of each tube region and area of the peripheral sheet region are then computed and incorporated into the analysis alongside the integrated fluorescence intensity in each tubule and sheet region. Multiple combinations of a peripheral sheet region and nearby tubules were selected in each individual image when possible and treated as distinct measurements of the local tubule radius.

Although the range of measurements for the tubule radius varied widely, the average over many structures in dozens of control wildtype cells (R = 51 +/-12 nm, mean +/-SD, n =141 independent regions) matched well to more detailed measurements reported with super-resolution imaging of tubular ER in COS-7 cells (R = 48 +/-9 nm^20^). Very similar tubule radii were measured in wildtype SH-SY5Y cells (R=51 +/-9 nm, n= 30). It should be noted that the current indirect approach has some advantages over the less accessible and more effort-intensive approaches. Namely, it can be carried out using standard confocal microscopy on live cells, thereby avoiding any potential artifacts associated with cell fixation.

A two-sample Kolmogorov-Smirnov test, designed for comparing data samples with no assumptions of Gaussian distribution, indicated that the distribution of tubule radii in cells with RTN4a overexpression or RTN4 knockout was significantly different from the distribution in control cells (*p* = 0.01 for COS-7 cells, *p* = 0.0003 for SH-SY5Y cells). Separate controls were carried out for each imaging run. Notably, when applied to comparing datasets from wildtype cells, the same statistical test gave non-significant *p*-values, giving no reason to reject the hypothesis that the two samples were drawn from the same distribution of tubule radii. Namely, a comparison of WT COS-7 cells with labeled calnexin vs Sec61β membrane protein yielded *p* = 0.7, and a comparison of WT COS-7 cells and WT SH-SY5Y cells (using fluorescently labeled Sec61β in both) yielded *p* = 0.5. These results imply that the measurements of tubule radii described here are relatively robust to the choice of visualized membrane protein, for the two cell types considered.

#### Photoactivation chase (iPAC)

The assay was performed using a confocal microscope (SP8, Leica, Wetzlar, Germany) with a controlled environment (37°C, 5% CO_2_). Images were acquired using a frame size of 512×512 pixels in the green (488nm excitation, 495-545nm emission), the red (555 nm excitation, 571-618 nm emission for TMR, or 567 nm excitation, 608-635 nm emission for mCherry), and the far-red (637 nm excitation, 645-700 nm emission) channels. After an acquisition of pre-photoactivation images, photoactivation illumination (405nm, 100% laser power for both Halotag-paJF646 and paGFP^ER^) was introduced in a region of interest (photoactivation spot) for a duration of 300 (COS-7) or 150 (SH-SY5Y) frames, using the Fly mode (enabling image recording during the photoconverting illumination) in the FRAP wizard. Intensity of the photoactivation channel at the photoactivation spot as well as other regions of interest were plotted as a function of time during photoactivation, and fitted to a double-exponential function, to extract the half time to reach plateau (t_1/2_), that was used as an estimator of Halotag-fused proteins or paGFP^ER^ mobility.

#### Photoconversion image analysis

The spread of proteins from the photoactivation site was quantified using custom code for image analysis and median rise-time estimation written in Matlab (version 9.5.0, R2018b, Natick, Massachusetts: The MathWorks Inc.). The cell and nucleus shape were manually segmented from images of ER luminal marker (mCherry^ER^). The segmented area, consisting of the region inside the cell boundary but outside the nucleus, was eroded by 1 micron to avoid edge effects.

The photoactivation region was directly segmented from the channel focused on the activation spot during the photoactivation phase. The centroid of the photoactivation region was set as the photoactivation centre. The photoactivation region, with a 1μm buffer, was removed from the cell area to be analysed.

The segmented cell area was first divided into concentric ring regions of width μm, with the inner radius starting at 1 um intervals in distance from the photoactivation center. Each ring was then divided into wedge-shaped ROIs of arc-length 2 um, shifted by 1 μm around the circle. Each wedge ROI was intersected with the segmented cell area, and only those ROIs with at least 2 μm^2^ of valid area (1 μm^2^ in SH-SY5Y cells) were retained for analysis.

Each of these ROIs yielded a fluorescence intensity time trace normalized by the ROI area (Fig. 2A, right).

For each cell, the intensity time series after the start of photoactivation was fitted to a double-exponential function rising from the pre-photoactivation value to a maximum value set by the final fluorescence of the photoactivated region. This fitting reflects the assumption that at infinite time, the fluorescence in the whole cell will equilibrate to that of the photoactivated region, even though the imaged timescales are generally too short to see this saturation for ROIs far from the photoactivated center. For the COS-7 RTN4a OE dataset (Fig. 2E) significant photobleaching was observed and an exponentially decaying photo-bleaching correction was included for both control and OE cells.

The double-exponential fits were used to compute a half-time for the signal rise. Curves with insufficient signal-to-noise ratio (signal range less than twice the pre-photoactivation standard deviation) were removed from the analysis, as were those giving a half-time more than 5 times the recorded timespan (∼70 seconds). The median half-time is computed for all remaining wedge regions at a given radius from the photoactivation centre, from all cells in each dataset. Error bars were obtained by bootstrapping individual cells from each dataset and repeating the analysis. The error bars therefore give an indication of the cell-to-cell variation in each reported half-time. The code is available on https://github.com/lenafabr/photoActivationAnalysis.git.

### Simulations of transport in ER tubular networks

Numerical simulations of intra-ER transport were carried out using a finite volume (FVM) approach, treating particle concentrations as continuous field variables defined on a network of one-dimensional edges connected at point-like junctions.

#### 1. Network geometry

Two-dimensional cell-shaped domains were extracted from images of ER luminal marker in 8 COS-7 cells (same cells used for membrane protein spreading analysis in Fig. 2B). The cytoplasmic domain was segmented as described in methods for photoactivation analysis. The domain was filled with a regular honeycomb network structure of edge length 1 μm, truncated at the domain boundaries.

Network edges were discretized into one-dimensional mesh cells of length δx, with δx < 0.1 μm. Each non-terminal junction node corresponded to a mesh cell of total length Σ δxi/2, where δxi is the mesh size on each of the adjacent edges.

#### 2. Simulation of photoactivated particle spreading

The photoactivated centre for each simulated cell was selected randomly, at a distance of 4-7 μm from the nucleus. The photoactivated region consisted of all mesh cells centred within 1.4 μm of the activation centre.

The concentration of photoactivated particles (c) is assumed to evolve according to the advection-diffusion equation

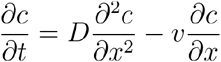

with c = 1 fixed within the photoactivation region, and with reflecting boundaries at all terminals (degree 1) nodes in the network structure. Initial conditions were set to c = 0 outside the photoactivated region. Simulations with diffusion only have v=0. Simulations including active flow set v = ±20 μm/s, with the flow direction selected randomly along each network edge. The flow direction on each edge remains persistent over a timescale of τ=0.1sec, with the flow reversal modeled as an independent Poisson process on each edge. At every time-step, for each edge, the flow direction is reversed with probability 1 - exp(-dt/τ).

The concentration field was evolved forward in discrete time-steps (dt = 5*10^−5^ sec), with forward Euler steps employed for the diffusive component (using a centered differencing method for the spatial differentiation) and a Lax-Wendroff scheme for the advective component. The choice of time step and mesh size satisfies the CFL criterion. Numerical convergence and stability were further established by testing that the resulting spatiotemporal concentration profiles were unchanged if either the mesh size or the timestep were decreased by a factor of two. Details of the finite volume numerical evolution scheme are provided together with our custom code (in Fortran 90; available on https://github.com/lenafabr/networkSpreadFVM.git).

The resulting concentration profiles on the meshed network were processed into images comparable to those collected for experimental data. Namely, the concentration field at each time interval of 0.14 sec was visualized (in Matlab, with a grayscale colormap) and pixelated into a 512 x 512 pixel image. The image was then blurred with a spatial Gaussian filter with σ=1 μm, and augmented with Gaussian noise with σ = 0.005 (concentration units). Resulting images were then processed to select wedge-shaped ROIs, trace ROI signals over time, and extract a characteristic half-time for the signal rise, exactly as for experimental data.

#### 3. Simulation of calcium dynamics

Our model for intra-ER calcium dynamics incorporates binding to calcium buffer proteins (with dissociation constant Kd), diffusion of free calcium ions (diffusivity Dc) and of buffer proteins (diffusivity Dp), and advective flow of velocity v. We consider the limit of fast calcium on and off rates, so that the calcium binding is assumed to be equilibrated at each time-step. The protein diffusivity is set to Dp = 2.8 μm^2^/s, based on our latest single particle tracking measurements in the ER lumen. Prior measurements of cytoplasmic diffusivity of calcium and calcium-binding proteins indicate that free calcium diffuses on the order of ten times faster compared to bound calcium ^34^. We therefore set the diffusivity of free calcium in the ER lumen to be a factor of ten higher than the buffer proteins (DC = 28 μm^2^/s). The dissociation constant is taken as KD = 0.3 mM, the initial free calcium concentration is set to be spatially uniform at 1 mM, and the initial concentration of buffer protein sites is set to 20 mM, in keeping with the weak binding strength and high calcium binding capacity of calreticulin proteins ^35^

The evolution of two fields is tracked over time and space: U(x,t) represents the concentration of unbound calcium ions and P(x,t) represents the total concentration of buffer protein sites (including bound and unbound). The time evolution of these fields is defined by

where B is the concentration of bound calcium ions, set at each time-step according to the equilibrium assumption: B = UP/(U+KD). The change in total protein (ΔP) and total calcium Δ(U+B) at each timestep is computed according to the usual finite

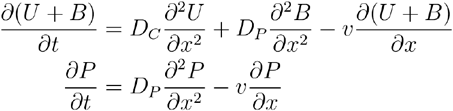

volume approach (Euler forward step with centred spatial differences for the diffusive components, Lax-Wendroff scheme for the advective component). The corresponding change in free calcium is then given by

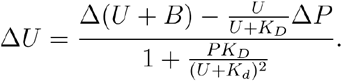

This approach explicitly maintains both equilibrium binding and mass conservation (in the absence of sources and sinks) over the time evolution.

This dynamic model is applied to an ER network extracted from a single confocal image of a COS-7 cells (one of the wildtype cells used for tubule width measurement in Fig. 1C). The image analysis was done using publicly available machine learning-based Ilastik software ^36^ for segmentation of a peripheral ER network structure, followed by the Matlab “skeletonize” function and custom-built code for grouping nearby nodes and tracing individual edges.

To model calcium release from the ER, we introduce a permeable region of radius 3μm and permeability p. For mesh cells within this region, a term of -pU*dt/δx is added to the equation for ΔU above, where δx is the mesh length. We make an upper limit estimate for the permeability coefficient by assuming that the tubule membrane within the permeable region is fully packed with calcium release channels, and that ions are free to move diffusively through the channel pores. Based on the published structure of the IP3R channel (PDB ID: 3JAV), the cross-sectional diameter of the channel cytoplasmic domain is approximately 12 nm, and the pore diameter is approximately 1 nm. The three-dimensional permeability of a section of ER tubule is then estimated according to p3D = DCA/h where h = 4 nm is the membrane thickness, A = (2π r) δx * (1nm)^2^/(12nm)^2^ is the total pore area in the mesh unit, and r = 50 nm is the tube radius. The permeability in the 1D model is then set to p = p3D/(πr^2^).

The dynamic system is evolved forward with timesteps of 2 * 10^−5^ sec, for a total simulation time of 20 sec. Random active velocities are represented as described for the photoactivated spread simulations, with 10 independent runs incorporating different realizations of the random velocity directions. Total calcium released from the permeable region is tracked over time by integrating the outward flux -pU * dt/δx at each time-step. Thick lines in Fig. 3A indicate the mean values over the 10 runs for each condition.

#### Photo-uncaging of caged-IP_3_ and imaging the ER Ca^2+^ release measurements

Prior to microscope imaging, cells were transfected to express GCaMP3^ER^, an ER membrane-tethered Ca^2+^ sensor, and treated with caged-IP_3_ (3 μM, 3 hours). For imaging, cells were transferred to the microscopy chamber of a confocal microscope (SP8, Leica, Wetzlar, Germany) with a controlled environment (37°C, 5% CO_2_). Images were acquired using a frame size of 512 × 512 pixels in the green channel (488 nm excitation, 510–530 nm emission). After an acquisition of pre-photo-uncaging images, photo-uncaging of caged-IP_3_ was achieved by illumination (405 nm, 20% laser power) in a region of interest (3 x 3 μm) for a duration of 300 frames using the Fly mode. Fluorescence intensity in the region of interest (F) was normalised to intensity at the beginning of photo-uncaging (F0) and plotted against time. The Area Under the Curve (AUC) was quantified for the duration of photo-uncaging to determine the amount of ER Ca^2+^ release.

#### Fluorescence Lifetime Imaging Microscopy (FLIM)

Prior to FLIM experiments, SH-SY5Y cells (WT or KO lines) were transfected to express ER-Crowdedness or D4ER probe (PMID: 27598166) for luminal calcium measurements and RTN4-SNAP for overexpression (see plasmid list for full details). Overexpression of RTN4a was monitored by labelling with SNAP-Cell 647-SiR. FLIM was carried out using a pulsed (80 MHz) wight light laser and HyDX or HyDS detectors in counting mode (STELLARIS8, Leica, Wetzlar, Germany). The excitation/emission was defined as 434/445-490 or 438/450-490 nm wavelength for D4ER and ER-crowding probes respectively. Images were acquired using a frame size of 512 × 512 pixels, and settings were set to reach 5000 photons/pixel. Images were processed using Leica STELLARIS8 FLIM wizard. ROIs were drawn around individual cells and data fitted with a monoexponential decay function. [Ca^2+^] FLIM analyses performed as in^37^. For D4ER, maximal calcium condition was determined as the ROI with minimal lifetime and minimal calcium condition (corresponding to maximum lifetime) was obtained by adding Thapsigargin (3 µM) to the culture media after baseline measurements.

#### Neurite outgrowth analyses

The outgrowth assays were obtained by automatic microscopy imaging (Incucyte, Satorius) of Human iPSCs. Cells were seeded on day 3 on a 96 well plate at a density of 20,000 cells per well, and from day 4 were repeatedly phase-imaged every 3 hours a 20x resolution. The resulting phase image stacks were subjected to the neurite detection module of the software accompanying the Incucyte microscope (Satorius) from which mask images of cell-body clusters and neurites were extracted. These images were then further processed through a custom Python code that 1. Computes the area covered by neurites and the number of different cell-body clusters and 2. Computes the neurite area and number of cell-body clusters for each well by adding the values obtained for the different field of view covering a well at each time point. The growth rates from Fig. 3E were obtained for each well by fitting a line to the normalised neurite area obtained by dividing the neurite area at each time point by the average area of cell-body clusters detected over all time points in this well.

### Generation of RTN4 knockout cell lines by CRISPR

The 20-nucleotide (nt) guide sequence in the guide RNA (gRNA) targeting *RTN4* was designed using the CRISPR Design tool (https://www.atum.bio/eCommerce/cas9/input). Oligonucleotides encompassing the guide sequence and BbsI restriction enzyme site overhangs (see Supplemental Table S2) were annealed, phosphorylated by polynucleotide kinase (PNK), and sub-cloned into the vector that expressed Cas9 from *Streptococcus pyogenes* together with a GFP, pSpCas9-2A-GFP (Addgene #48138, Cambridge, MA, USA). 24 hours post-transfection, gRNA and Cas9 expressing cells were sorted by cell sorter (FACS Melody, BD Biosciences, San Jose, CA) (SH-SY5Y and COS-7), or sorted manually (iPSCs). Each single clone was subjected to genomic DNA extraction and PCR to identify insertions and deletions (indels) by non-homologous end-joining (NHEJ). PCR products were subsequently sequenced using forward primers used in the PCR. Selected clones were subsequently subjected to Western blot to confirm knockout at the protein level. Primer information is shown in the Supplemental Table S2.

### Protein expression analyses

The cells were lysed by radioimmunoprecipitation assay (RIPA) buffer and protein concentrations of lysate were determined using a BCA Protein Assay Kit (#23225, Thermo Fisher Scientific Inc., Waltham, MA, USA). Equivalent amounts of proteins were subjected to sodium dodecyl sulfate-polyacrylamide gel electrophoresis (SDS-PAGE) and transferred to poly-vinylidenefluoride (PVDF) membrane. The membranes were blocked and incubated with primary antibodies to RTN4 and beta-tubulin at 1:1000 each. Fluorescent dye-conjugated secondary antibodies were used at 1:3000 and the proteins were detected using ChemiDoc MP Imaging System (Bio-Rad, Hercules, CA, USA).

### Statistics and reproducibility

Statistical analyses and visualisation were performed using Prizm 9. Error bars, P values and statistical tests and sample sizes are reported in the figure legends. All experiments were performed independently at least three times. Statistical differences between probability distributions were assessed using two-way Kolmogorov–Smirnov tests and statistical differences between distribution medians were assessed using two-sided Mann–Whitney U-tests.

**Fig S1.**
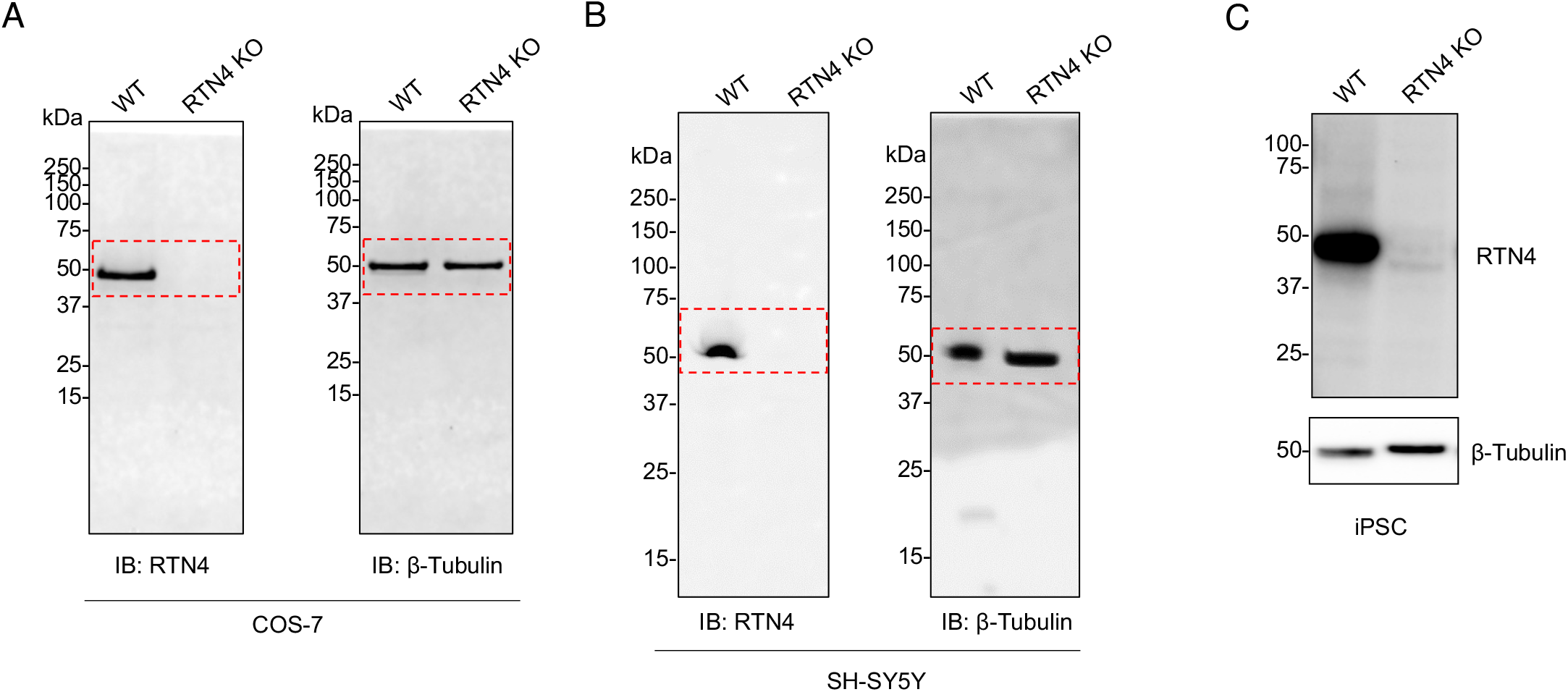
Western blot analyses of CRSPR RTN4 knockout cell lines. Full membranes of blots in A. COS-7, B. SH-SY5Y (Fig. 1D) and C. undifferentiated iNeuron (iPS. Fig. 4) cells.

**Video 1. Exclusion of luminal content by endogenous RTN4 in fixed COS-7 cell**. A fluorescent micrograph from Fig. 1A of immunolabeled endogenous RTN4 (yellow) and exogenousely expressed mCherry^ER^ (magenta).

**Video 2. paGFP**^**ER**^ **photoactivation pulse chase**. Note paGFP^ER^ (yellow) appearance and spread in response to local laser irradiation (405 nm). The ER network structure was visualised by mCherry^ER^ (magenta). Scale bar, 5 μm.

**Video 3. A simulation model of ER luminal transport**. Particle concentration is fixed within a localised region of radius 1.4 μm, representing the photoactivated area. Concentration over time is shown on a hexagonal lattice confined within a cell-shaped domain. Top: diffusive spreading only (with D = 3 μm^2^/s). Bottom: diffusion and flow on an active network, with randomly directly flow velocities of v = 20 μm/s, and persistence time 0.1 s on each edge. Right: Corresponding pixelated images of simulation snapshots, blurred over a 1 μm length scale and with added noise. These images were used to quantify signal rise times in individual wedge ROIs, analogous to the analysis of experimental iPAC data. Scale bars are 10 μm.

**Fig S2.**
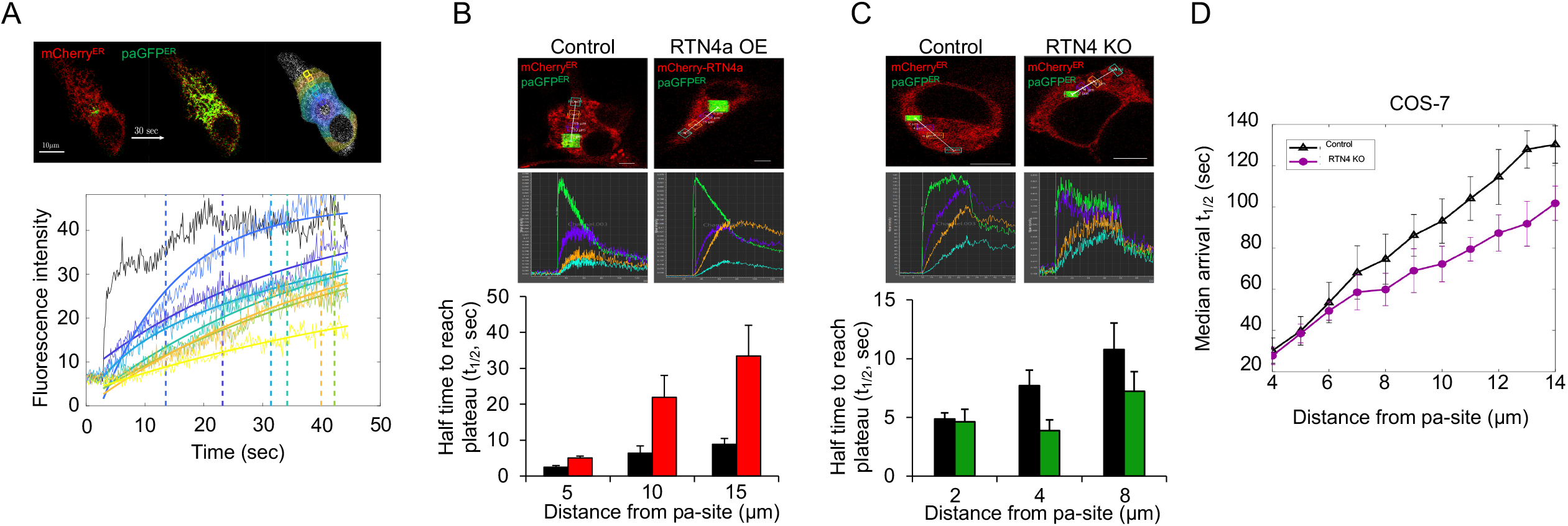
Additional analysis of paGFP^ER^ photoactivation pulse-chase assay. A. Representative raw images showing automated detection of paGFP^ER^ transport corresponding to Fig 2B, D, E, and F. B. Raw images of representative samples (Control and RTN4a OE, corresponding to Fig. 2E). Fluorescence intensity change over time was measured at manually selected ROIs with different distances from the photoactivation spot. Raw data during photoactivation period was fitted to a mono-exponential equation, and the half time to plateau (t_1/2_) was determined for each distance. C. Analyses as in B in this case comparing WT and RTN4 KO cells (corresponding to Fig. 2F). D. iPAC analysis as in Fig. 2A of RTN4 KO COS-7 cells. Note, control signal rise times match well to luminal protein data in Fig. 2B. RTN4 KO data results in faster signal spread to long distances, as observed for SH-SY5Y cells in Fig. 2F.

**Fig S3.**
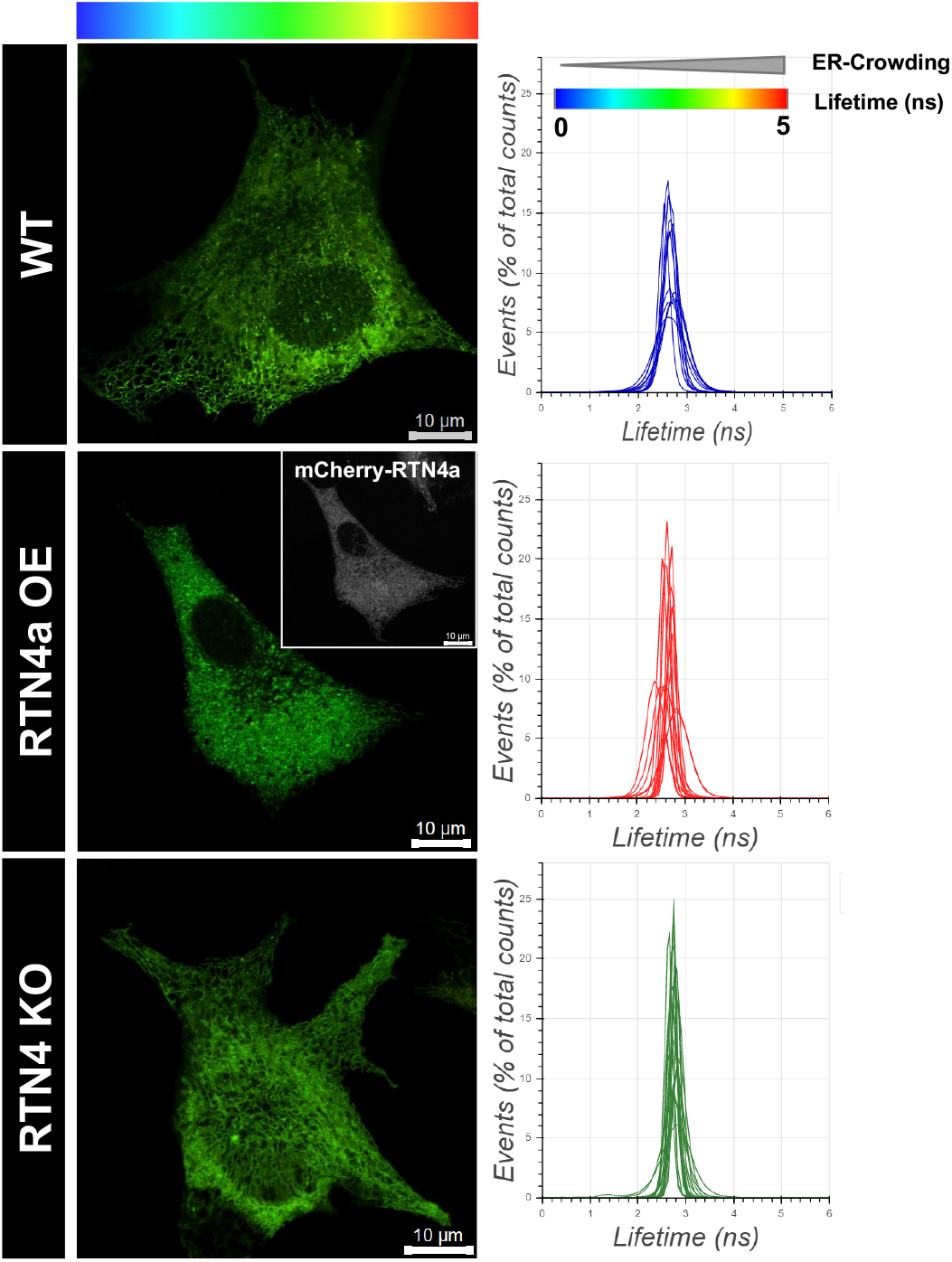
Source images for Fluorescence Lifetime measurements of ER crowdedness in Fig. 2G. Histograms show frequency distribution of lifetime values in their corresponding images.

**Fig S4.**
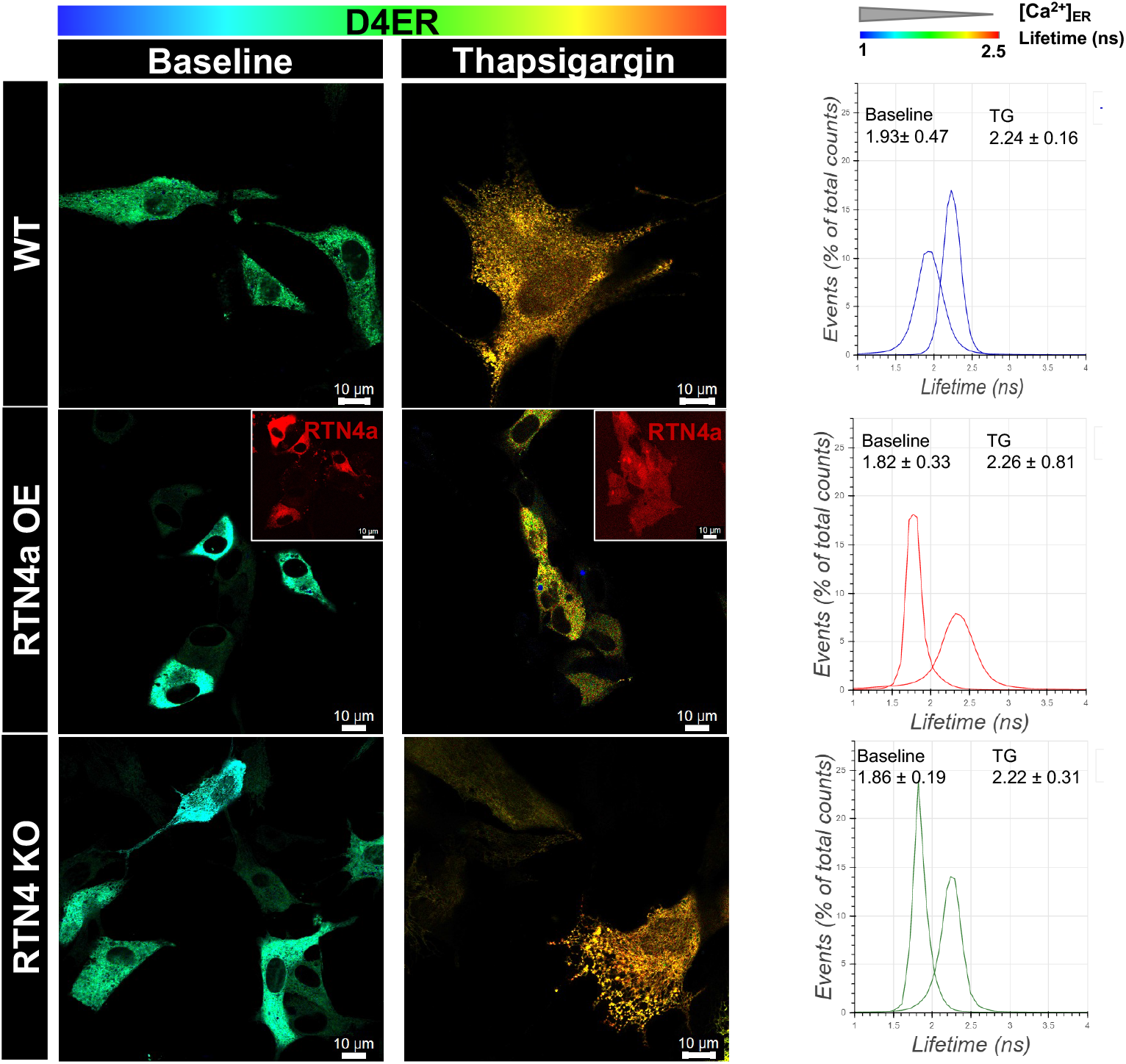
Source images for Fluorescence Lifetime measurements of ER Ca^2+^ of Fig. 3F. Shown are colour-coded FLIM images of D4ER lifetime at the baseline and after ER Ca^2+^ depletion by Thapsigargin treatment. Histograms show the frequency distribution (and the corresponding mean ± STD) of lifetime values in representative cells for each treatment condition.

**Video 4. A simulation of Ca**^**2+**^ **transport in the ER**. Ca^2+^ concentration profiles in an ER network structure, with a local permeabilised region (green circle) are shown over time. Simulations were run as in Fig.3A Left: diffusive transport only (D_p_ = 2.8 μm^2^/s, D_c_ = 28 μm^2^/s). Middle: Advective flows with v = 20 μm/s and persistence time 0.1 sec are included as well as diffusion. Right: cumulative calcium release from the permeable region, for the two simulation runs show.

**Video 5. Neuro-physiological Ca**^**2+**^ **fluctuation in iNeurons as in Fig 4**.

**Fig S3.**
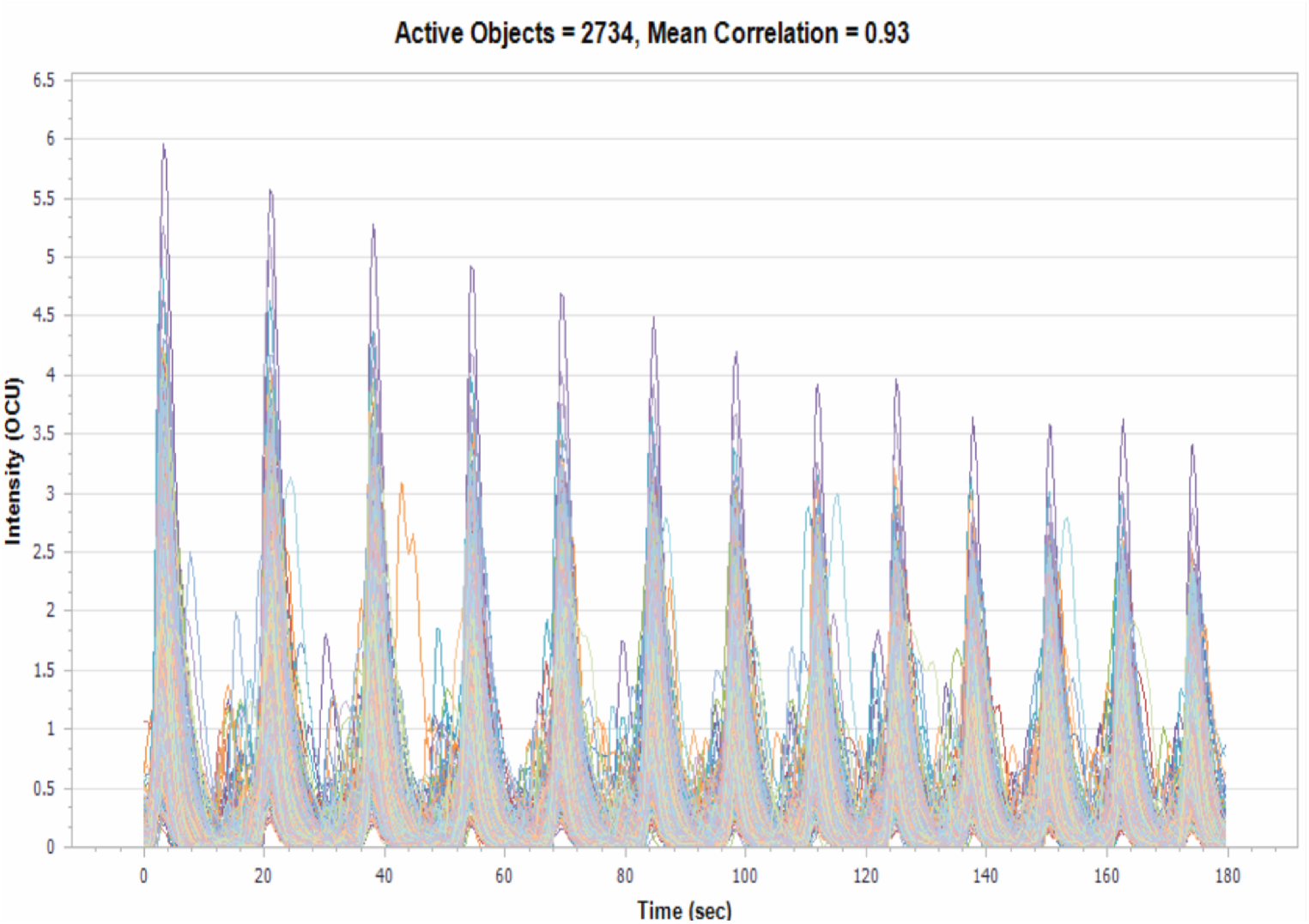
Spontaneous Ca^2+^ fluctuation in differentiated iNeurons. Neuronal Ca^2+^ fluctuation acquired as in Video 6 was analysed by the Incucyte S3 Neuronal Activity Analysis Software Module. Active objects indicate the number of objects (cells/cell clusters) that burst at least once above the minimum burst threshold over the total scan time. Mean correlation indicates neural network connectivity obtained by a comparison of every active object to every other object. 0 being completely random and 1 being highly synchronised.

**Video 6. Time-lapse phase contrast micrographs of neurite outgrowth in differentiated iNeurons as in Fig. 4D**.

**Fig S6.**
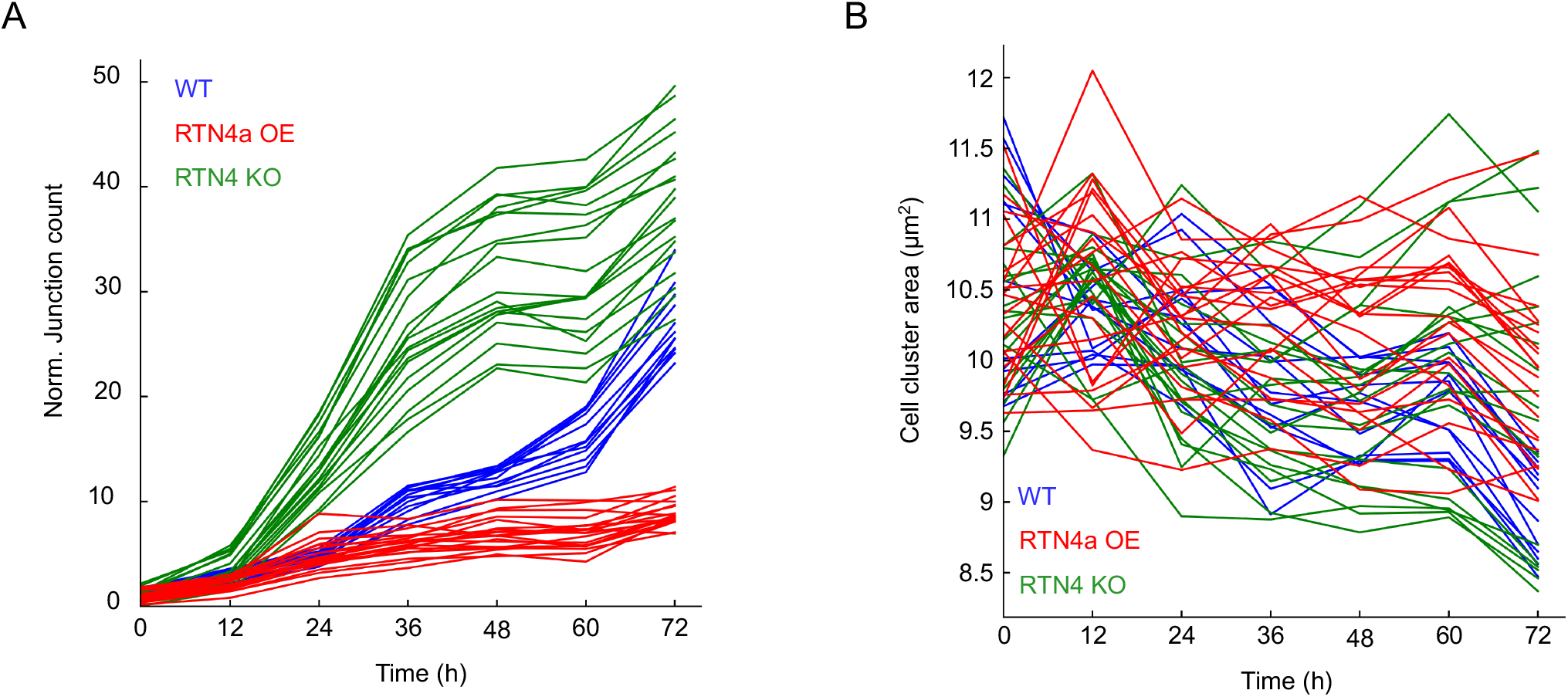
Quantification of neurite outgrowth. Further quantification of neurite outgrowth pattern. (A) Normalised number of junctions divided by the average number of cell body-clusters in the field of view as a function of time. B. Area of detected cell-body clusters as a function of time.

**Video 7. Neurite regeneration assay in iNeurons corresponding to Fig 4G and H**.

Plasmid list
| ID | Plasmid name | Description | Reference | First appearance | Label in figure |
| --- | --- | --- | --- | --- | --- |
| 1 | pSpCas9(BB)-2A-GFP | Cas9 from *S. pyogenes* with 2A-EGFP, and cloning backbone for sgRNA | Addgene #48138 | Materials and Methods | - |
| 2 | pSpCas9(BB)-2A-GFP_gRNA RTN4 | CRISPR for human reticulon 4 knockout | this study | Materials and Methods | - |
| 3 | pLV_Exp_Puro-TRE3G_XbaI_Hu Rtn3A-TEV-HaloTag | Mammalian expression of human Halo-RTN3A | this study | Figure 2 | RTN3A |
| 16 | pcDNA3-ER-GCaMP3 | Mammalian expression, ER-targeted Ca<sup>2+</sup> indicator GCaMP3; targeted to cytosolic face of the ER membrane. | PMID:25056880<br>Addgene #64854 | Figure 3 | GCaMP3<sup>ER</sup> |
| 32 | pFLAG_ER mCherry | Mammalian expression, ER-localized mCherry | PMID: 25073928 | Figure 1 | mCherry<sup>ER</sup> |
| 34 | Halo-Sec61b | Mammalian expression of Halo-Sec61b | PMID: 32875078 | Figure 1 | Halo-Sec61β |
| 55 | pmPA-GFP-ER-5 | Mammalian expression of ER-targeted photoactivatable GFP | Addgene #57132 | Figure 2 | paGFP<sup>ER</sup> |
| 57 | pMD1g/pRRE | 3rd generation lentiviral packaging plasmid; Contains Gag and Pol | Addgene#12251 | Figure 4 | - |
| 58 | pRSV-Rev | 3rd generation lentiviral packaging plasmid; Contains Rev | Addgene#12253 | Figure 4 | - |
| 59 | pDM2.G | VSVG envelope expressing plasmid for lentivirus | Addgene#12259 | Figure 4 | - |
| 88 | mCherry_SBP_RTN4A | Mammalian expression of mCherry-humanRTN4a | PMID: 30772082 | Figure 1 | RTN4a |
| 118 | ER-Crowding_FRET1_CRH2_KDEL_pCDNA3-1(+) | ER crowdeness plasmid | PMID: 30224760 | Figure 2 | ER-localised molecular crowding probe |
| 170 | pLV_Rtn4-SNAP | Mammalian expression of SNAP-humanRTN4a | this study | Figure 1 | SNAP-RTN4a |
| 188 | pLV_TRE3G_hRTN4a-Halo | Mammalian expression of human RTN4a-Halo | this study | Figure 4 | RTN4a-Halo |
| 220 | D4ER | FRET-based calcium sensor located in the ER lumen | PMID: 27598166 | Figure 3 | D4ER |
| 246 | Emerald-Calnexin | Mammalian expression of Emerald-Calnexin | Addgene #54021 | Figure 1 | - |
| 247 | mcherry_RTN4a | Mammalian expression of mCherry-humanRTN4a | PMID: 20955502 | Figure 1 | RTN4a |

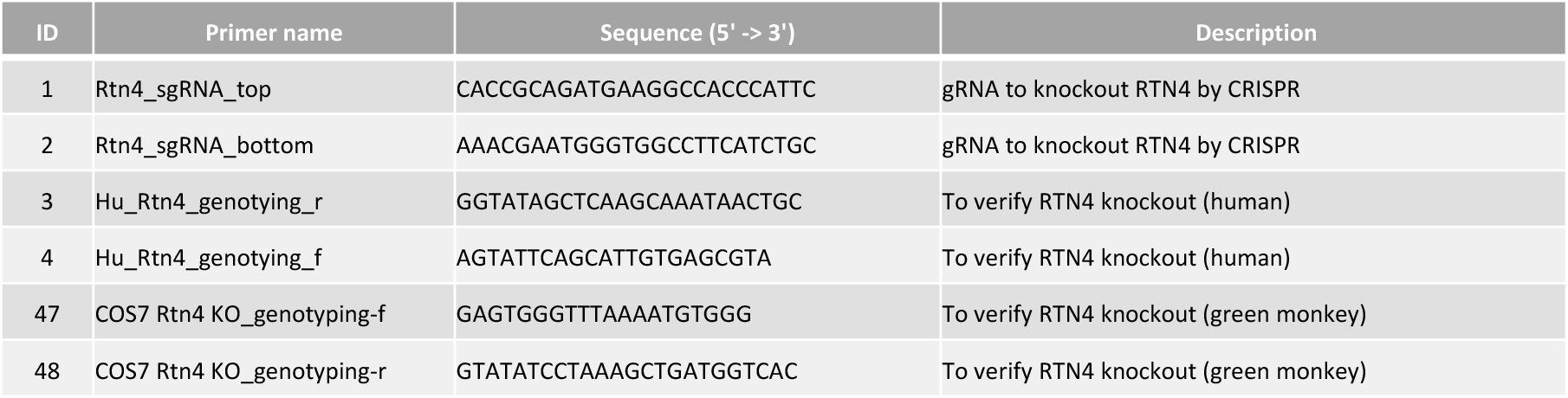

