## Supplementary figures and images for "Endoplasmic Reticulum morphological regulation by RTN4/NOGO modulates neuronal regeneration by curbing luminal transport"

### Video 1

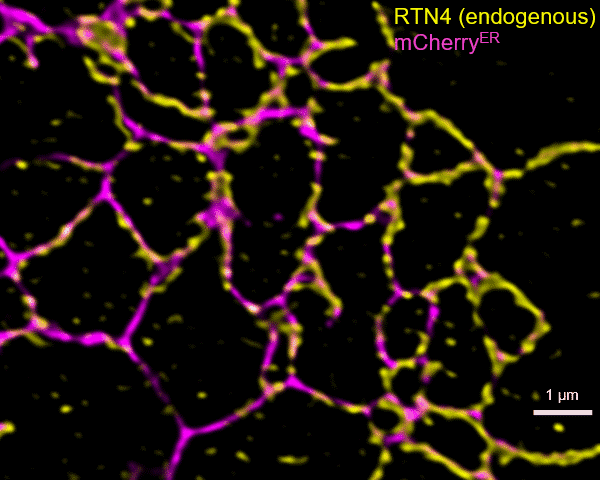
